# ENKEFALOS: Electrophysiological NetworKs Elicited From ALtered OmicS

**DOI:** 10.64898/2026.11.27.769720

**Authors:** Krish Upadhyay, Dylan Ukasik, Douglas Craig, Alan Dombkowski

**Affiliations:** Cranbrook Kingswood High School, Bloomfield Hills, MI 48304; Translational Neurosciences Program, Wayne State University, Detroit, MI 48202; Department of Computer Science, Wayne State University, Detroit, MI 48202; Department of Pediatrics, Wayne State University, Detroit, MI 48202

**Keywords:** electrophysiology, neuron, pathway, proteome, transcriptome

## Abstract

‘Omics’ analyses such as genomics, transcriptomics, and proteomics have proven to be critical in uncovering the cellular and physiological mechanisms of a wide range of diseases, including neurological disorders. These technologies often produce lists of thousands of differentially expressed (DE) genes or proteins. Despite the vast amount of information provided by these methods, their value is highly dependent upon the ability to interpret results within the appropriate biological context. Numerous bioinformatics tools are available to interrogate DE genes and identify their association to general cellular pathways, but a critical gap in neurological studies is a lack of computational tools to decipher DE genes specifically for their role in neuronal electrophysiology. This ability is essential for the effective utilization of omics data in studies of epilepsy and neurodegenerative diseases. To address this gap, we developed an easy to use, web-based bioinformatics tool entitled ENKEFALOS: Electrophysiological NetworKs Elicited From ALtered OmicS. Users can submit lists of DE genes or proteins to ENKEFALOS which identifies the subset that have a significant association with neuronal electrophysiological properties, based on publicly available patch clamp and RNA-seq data. ENKEFALOS then interrogates the subset of genes/proteins to identify network interactions, enriched cellular pathways, and key nodes. We demonstrate the utility of ENKEFALOS using a dataset of DE genes from a seminal study of epilepsy in tuberous sclerosis complex patients. ENKEFALOS is accessible at: https://appyters.maayanlab.cloud/#/.

**NEW & NOTEWORTHY:** We present a web-accessible, easy to use, and innovative bioinformatics tool that interprets lists of differentially expressed genes or proteins for their functional relevance to neuronal electrophysiology. ENKEFALOS (Electrophysiological NetworKs Elicited From ALtered OmicS) provides a means to link differentially expressed genes/proteins to changes in neuronal electrophysiological properties and highlight the underlying cellular pathways. Interactome networks and key signaling nodes are identified to enable discovery of new therapeutic targets in neurological disorders.

## INTRODUCTION

The primary strength of transcriptomic and proteomic technologies is the ability to concurrently quantify molecular expression for thousands of genes or proteins. The resultant genome-wide snapshot of expression is analogous to thousands of separate puzzle pieces scattered on a table. Their assembly into a comprehensive picture of cellular function requires the use of bioinformatics tools to categorize and assemble puzzle pieces (genes/proteins) into cellular components such as signaling pathways, functional units, and interactome networks [1-4]. This process is akin to finding all puzzle pieces that comprise elements of the scene, such as a barn or the trees in a landscape. Many excellent bioinformatics tools have been developed for this purpose [5-7].

The study of the cellular origins of neurological diseases is often focused on disruptions of neuronal electrophysiology. This is apparent in studies of epilepsy, but is also foundational to multiple sclerosis, Alzheimer’s disease, Parkinson’s disease, and other neurodegenerative disorders [8-10]. A critical gap in the use of functional “omics” technologies to further our understanding of neurological disorders is a lack of computational tools to decipher lengthy lists of genes or proteins for their role in neuronal electrophysiology.

To the best of our knowledge, there are currently no publicly available bioinformatics tools that are tailored to perform pathway and network analysis of lists of genes, specifically for their potential role in alterations of neuronal electrophysiology. To address this gap we developed a bioinformatics tool entitled ENKEFALOS: Electrophysiological NetworKs Elicited From ALtered OmicS. Importantly, ENKEFALOS was designed to be web-accessible, easy to use, and require little or no coding skill.

As the foundation for ENKEFALOS, we utilized previous published research that integrated publicly accessible datasets for gene expression and neuronal electrophysiology derived from patch clamp experiments [11]. That important work provided a means to examine relationships between gene expression and electrophysiological properties on an individual gene basis. Building on their groundwork, we devised a tool for omics-wide analyses that takes as input a list of DE genes or proteins, identifies the subset having expression profiles significantly correlated to neuronal electrophysiological properties, and performs analysis of pathways and interactome networks associated with the identified genes. This approach allows an investigator to determine the collective effect on electrophysiology from numerous differentially expressed genes or proteins. Furthermore, ENKEFALOS enables identification of key nodes (genes/proteins) that are highly interconnected in the identified networks and thus represent likely drivers of pathogenesis and potential therapeutic targets.

To demonstrate how ENKEFALOS can uncover altered genes, networks, and key nodes that are involved in neuronal electrophysiology, we used ENKEFALOS to interrogate a published dataset from a seminal study of epilepsy in tuberous sclerosis complex (TSC). ENKEFALOS uncovered key pathways and nodes linked to alterations of neuronal electrophysiology that are supported by recent literature and warrant further investigation for their role in TSC epilepsy. ENKEFALOS has potential application in a wide range of neurological studies that employ omics analyses.

## MATERIALS AND METHODS

### ENKEFALOS architecture

An overview of the ENKEFALOS architecture is provided in **Figure 1**. To facilitate access by the broad research community, we created ENKEFALOS to run under the Appyter framework, thus enabling web-based access via a simplified user interface that does not require programming experience [12]. ENKEFALOS is accessed from the Appyter catalog (https://appyters.maayanlab.cloud/#/) and can be run either on the Appyter server or on the user’s local computer. The primary user input is a list of gene or protein symbols which can be pasted into the user interface or uploaded from a file. Input genes or proteins are queried against a database that links neuronal gene expression to electrophysiological properties. Output of ENKEFALOS includes a list of the subset of input genes that have a statistically significant association between their expression and electrophysiological properties. Additional output includes: an interaction network of the significant genes/proteins, key nodes of the network based on the level of connectivity, and enriched biological pathways. A user guide is available as Supplemental File S1 and at the GitHub site. The main components of the application are described in the subsections below.

**Figure 1.**
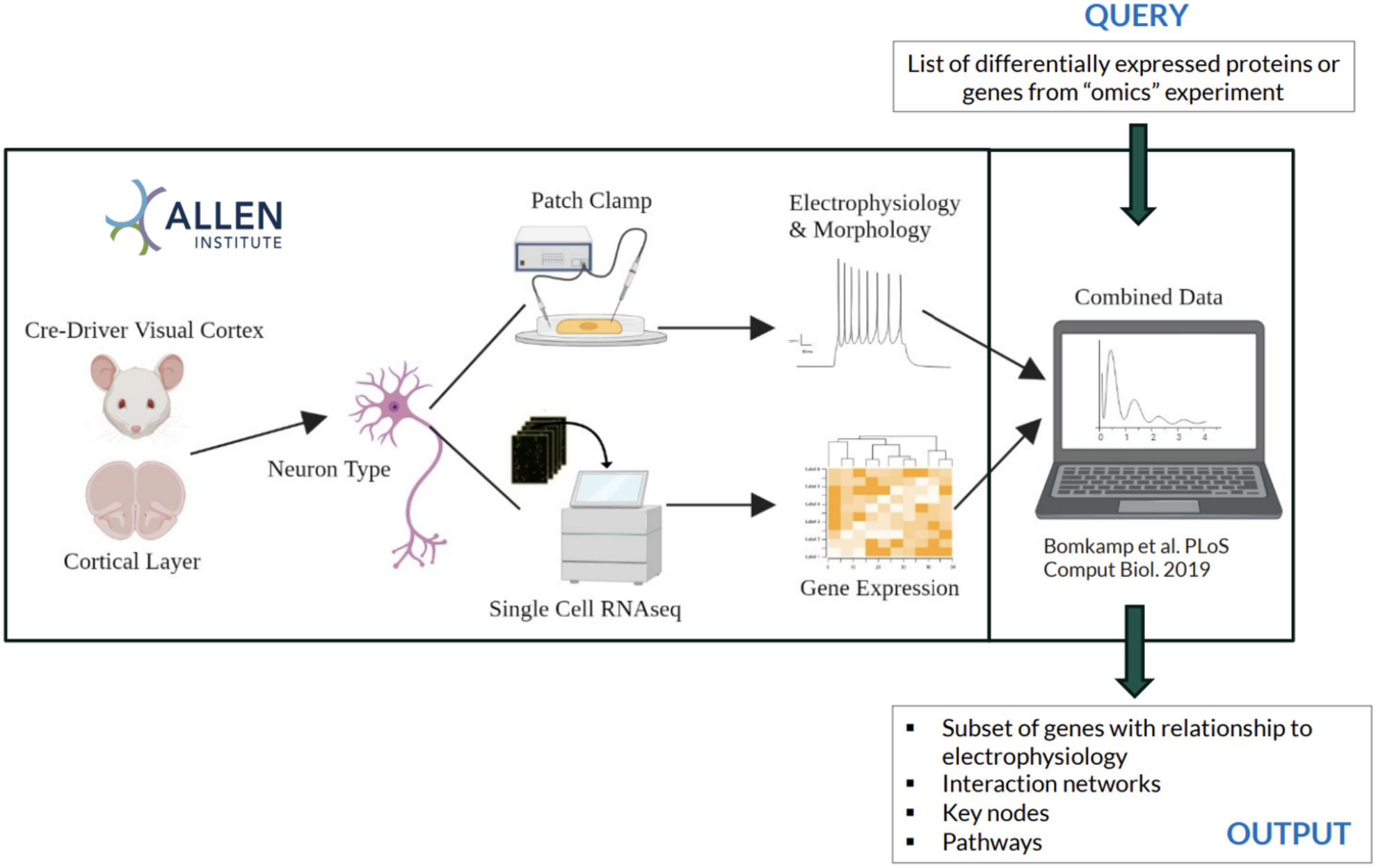
Enkefalos architecture. Transcriptomic and patch clamp data from the Allen Institute was previously assembled into a database by Bomkamp et al. to link individual genes to electrophysiological properties. We built Enkefalos around the database to enable queries using large sets of genes or proteins and provide interrogation of results using pathway and network analyses.

### Python

ENKEFALOS was created using the Python 3 programming language. All code and data are available at Github (https://github.com/MaayanLab/appyter-catalog/tree/main/appyters/ENKEFALOS_1/; https://github.com/MaayanLab/appyter-catalog/tree/main/appyters/ENKEFALOS_2). The Pandas library in Python was used for data analyses. Underlying data for gene expression and electrophysiological/morphological properties are derived from the Pavlidis study and were formatted into Excel spreadsheets [11]. Pandas creates a data frame for each spreadsheet which are then queried with user input. Additional data frames are created as the program progresses to record significant correlations and other specificities.

### Electrophysiological and transcriptomic data

The underlying electrophysiological, morphological, and omics datasets used by ENKEFALOS were derived from a transcriptomic correlates study previously published by the Pavlidis Lab [11].Their dataset was obtained from publicly available single-cell transcriptomic and patch-clamp data at the Allen Institute of Brain Sciences (https://celltypes.brain-map.org/). The Allen Institute used Cre-driven reporters to select subtypes of inhibitory (GABAergic) and excitatory (glutamatergic) neurons from the primary visual cortex of mice. Selected cells were subject to RNA-seq and whole cell patch-clamp experiments, providing transcriptomic and electrophysiological data for each cell type. The Pavlidis lab integrated these data for correlative analysis and identification of “class-driven” relationships between gene expression and electrophysiological properties, i.e. differential between excitatory and inhibitory neurons, and non-class driven relationships within a neuronal class. Additional details are available in the associated paper from the Pavlidis laboratory [11]. While the underlying data is derived from murine experiments, it is expected that electrophysiological associations derived from mice are applicable to other mammalian species, including human. However, some input human genes may not match to mouse orthologs.

Electrophysiological properties include: action potential threshold, rheobase, adaptation ratio, input-output curve slope, max firing frequency, afterhyperpolarization threshold, interspike interval analyses, sag, input resistance, resting membrane potential, time constant tau, afterhyperpolarization amplitude, average interspike interval, afterhyperpolarization half-width, capacitance, and latency (Supplemental File S2).

### Interactome networks

Gene/protein interaction networks and functional enrichment analyses derived within ENKEFALOS are extracted from the STRING database [6]. Using the STRING API, ENKEFALOS obtains a link to a STRING webpage displaying the network derived from the user’s genes/proteins of interest. Functional enrichment analyses are performed by STRING using other databases, including but not limited to KEGG pathways, Reactome, Gene Ontology Consortium, and PubMed. The NCBI species identifier code for humans is used in ENKEFALOS queries.

The NetworkX library (networkx.org) was used for network analyses of STRING interactome results. After STRING analysis, each node in the gene/protein interactome was tabulated using a NetworkX algorithm to count the number of interactors each gene/protein has in the network. Moreover, the creation of subset networks was performed in NetworkX, which are passed back to STRING again for subsequent functional enrichment analyses.

### Appyter

ENKEFALOS is accessible under the Appyter catalog developed by the Ma’ayan lab (https://appyters.maayanlab.cloud/#/ENKEFALOS_1, https://appyters.maayanlab.cloud/#/ENKEFALOS_2). ENKEFALOS was first converted into a .ipynb file to make it compatible with Jupyter, primarily for the purpose of having markdowns included in the ENKEFALOS code and for the code to gain the ability to be executed in subsections, rather than as a single script. Appyter magics were then implemented into the existing .ipynb files, thus providing an entry form that enables users to upload their data and selectable parameters for a multitude of data analysis workflows [12]. Once the inputs are submitted, the Appyter executes the corresponding notebook in the cloud, producing visual and downloadable outputs from the ENKEFALOS analyses without requiring the user to interact directly with the code. ENKEFALOS was crafted into two Appyters: one being the primary gene/protein-list based tool, while the other Appyter is secondary and dependent on analyses completed by the primary Appyter.

### Parameter selection and output

The user can specify a false-discovery rate cutoff (FDR, q-value) for selection of genes based on the significance level of the correlation between gene expression and electrophysiological properties. The q-value is extracted from the Pavlidis Lab database [11]. Additionally, users can select the method for modeling gene-property relationships with respect to cell class, as was defined by the Pavlidis group. Options are: 1) the class-conditional model (CC) which considers cell class (excitatory or inhibitory) in modelling the relationship between gene and property; or 2) class-independent model (CI) which ignores cell class in the model; or 3) interaction model (Int) which identifies gene-property relationships that differ between the two classes.

Gene-property relationships that meet the statistical cutoff under the specified cell class model are output in a list with the q-value under “Significant Correlations.” A CSV file of the selected list can be downloaded. Note that a gene may be listed multiple times, reflecting multiple electrophysiological properties. A non-redundant list of genes is shown under “Genes with Significant Interactions.”

From the subset of input genes/proteins having a significant correlation to electrophysiological properties, ENKEFALOS identifies associated biological pathways, interaction networks, and key nodes. A network map is displayed and shows how each of the genes/proteins interact with each other. ENKEFALOS further tabulates the number of interactions for each node (gene/protein) and provides a downloadable table. Highly connected nodes are potential key drivers of altered electrophysiology. The output from STRING also includes enriched biological pathways and ontologies. Readers are encouraged to use the help tab in STRING to facilitate use of this resource.

### Secondary analysis

Additional analyses are available in ENKEFALOS to characterize specific genes identified in the primary output, for example, a highly interconnected key node. These features are further demonstrated in the case study. The Appyter button labeled “Secondary Analysis” allows the user to specify a gene of focus, select an electrophysiological property that was linked to the gene in the primary analysis, and upload the list of significant genes obtained from the primary analysis. ENKEFALOS will then output a scatter plot, derived from the Pavladis dataset, that shows the correlation between the select gene and the specified electrophysiological property. Data from inhibitory and excitatory neurons are shown. The underlying data used to create the scatter plot can be downloaded as a CSV file. To explore the direct interactome of a specified gene/protein, ENKEFALOS creates a STRING network plot of the subset of genes from the primary analysis that directly interact with the specified gene of interest, rather than through secondary nodes. This feature enables the user to further investigate the direct interactome network and pathways associated with a potential key node.

## RESULTS

### Case study: Epilepsy in tuberous sclerosis complex

To demonstrate the utility of ENKEFALOS we analyzed gene expression data from a seminal and highly cited transcriptomics study that investigated gene expression in epilepsy of tuberous sclerosis complex (TSC). TSC is an autosomal dominant genetic disorder that arises from mutations in the TSC1 and/or TSC2 genes and leads to the formation of benign tumors in multiple organs, including cortical tubers in the brain [13, 14]. Approximately 60% of TSC patients suffer from drug resistant epilepsy. Tubers are often identified as seizure foci in TSC, but the exact mechanism of seizure induction has yet to be elucidated. Understanding the electrophysiological properties of DE genes in epileptogenic TSC tissue is critical to identifying potential mechanisms of seizure induction.

Using gene expression microarrays, the Aronica laboratory performed a seminal transcriptomic study of TSC tubers that were surgically resected to treat drug-resistant epilepsy [15]. They compared epileptogenic tubers with normal perituberal tissue and autopsy controls. The authors identified many DE genes and noted the genes represent a mixture of those potentially involved in tuber formation as well as seizure induction. However, it is important to disentangle the transcriptomic alterations associated with epileptogenesis from those associated with lesion (tuber) formation to better understand the molecular events that drive seizures. Here, to demonstrate focus on epileptogenesis we used ENKEFALOS to identify the subset of DE genes involved in neuronal electrophysiology alterations, along with their interactome networks and signaling pathways.

To create our test set of genes from the Aronica study, we used the GEO2R tool at the NCBI Gene Expression Omnibus and dataset GSE16969 of Affymetrix data. We applied a moderated t-test to compare gene expression in epileptogenic tuber tissue (4) to autopsy controls (4). We selected transcript probes with significantly increased expression in epileptogenic tissue with a false discovery rate (FDR) ≤ 10%, resulting in 6164 significant probes. The fold change values for redundant probes of the same gene were averaged, providing 3976 unique genes (Supplemental File S3). These genes were input into ENKEFALOS, which identified 7286 significant associations to electrophysiological properties (FDR ≤ 10%). Note that an individual gene may be significantly correlated to multiple electrophysiological properties. Of these, ENKEFALOS identified 1229 unique genes

The integrated pathway and network analysis of ENKEFALOS identified signal transduction as the most significantly enriched Reactome pathway (FDR= 2.27e^-9^). Among specific signal transduction pathways was receptor tyrosine kinase (RTK) signaling with 57 genes (FDR= 0.0168). Analysis via STRING of the 57 RTK genes reveals the most significant sub-pathway was EGF/EGFR signaling (Wikipathways, FDR=1.94^e-11^). The key node analysis feature in ENKEFALOS further highlights the importance of EGFR signaling as detailed below.

ENKEFALOS tabulates connectivity for each node (gene or protein) among the network of genes having significant electrophysiological correlation. Highly connected genes are likely to be key nodes in the disease or biological process being studied. The 20 most highly connected genes are shown in **Table 1**, and EGFR was the second most highly connected node. Using these 20 genes we performed a secondary pathway analysis within STRING. The most significant Reactome pathways associated with the 20 genes were: 1) Signal transduction, FDR=4.02^e-8^; 2) Diseases of signal transduction by growth factor receptors and second messengers, FDR=1.91^e-7^; 3) Disease, FDR=5.14^e-7^; and 4) Signaling by receptor tyrosine kinases, FDR=1.38^e-5^. A Sankey diagram in Figure 2A illustrates the hierarchical relationship of the 20 genes to these biological pathways. The interactome of the 20 key node genes is shown in Figure 2B. The five most highly connected genes are MYC, EGFR, HDAC1, JUN, and STAT3, and they form an interconnected supernode (Figure 2C).

**Table 1.** The 20 most highly connected nodes identified by ENKEFALOS in our case study of induced genes in epilepsy of tuberous sclerosis complex. Each gene has a statistically significant relationship to a neuronal electrophysiological property and is highly interconnected in network analysis.

| Gene | Interactions |
|---|---|
| MYC | 135 |
| EGFR | 123 |
| HDAC1 | 97 |
| JUN | 94 |
| STAT3 | 78 |
| CASP3 | 73 |
| VCL | 61 |
| SOX2 | 58 |
| LMNA | 57 |
| POLR2A | 56 |
| SMAD2 | 54 |
| ITGB1 | 53 |
| TJP1 | 52 |
| EIF4E | 52 |
| RUNX2 | 51 |
| PRKCA | 50 |
| IQGAP1 | 50 |
| DICER1 | 49 |
| GART | 48 |
| CHD4 | 47 |

**Figure 2.**
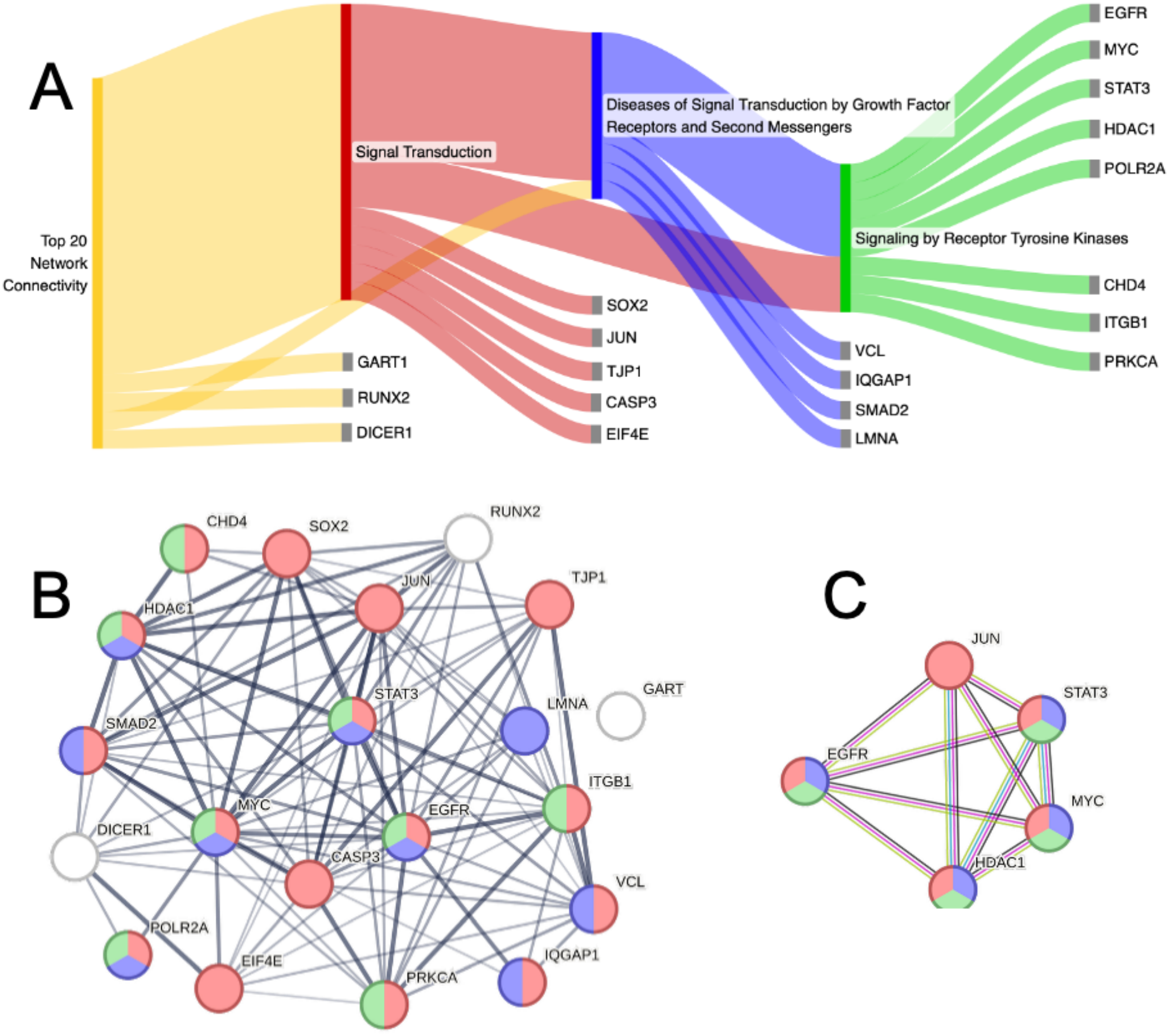
Enkefalos analysis of differentially expressed genes from a seminal transcriptomic study of TSC brain tissue surgically resected to treat epilepsy. Enkefalos identified 20 highly interconnected genes linked to neuronal electrophysiology and reported associated cellular pathways. **A)** Sankey diagram of significant Reactome pathways of the 20 highly interconnected genes reveals enrichment for signal transduction. **B)** The interactome of the 20 genes, as determined by STRING. **C)** The five most highly connected genes form a supernode centered on EGFR signalling.

Interestingly, ENKEFALOS revealed that EGFR gene expression is significantly correlated with several electrophysiological measures within inhibitory neurons but not excitatory neurons. An example is afterhyperpolarization amplitude (AHP) which is anticorrelated with EGFR expression in inhibitory neurons (**Figure 3**). A significant correlation was not observed in excitatory neurons. The AHP modulates the rate of subsequent firing.

**Figure 3.**
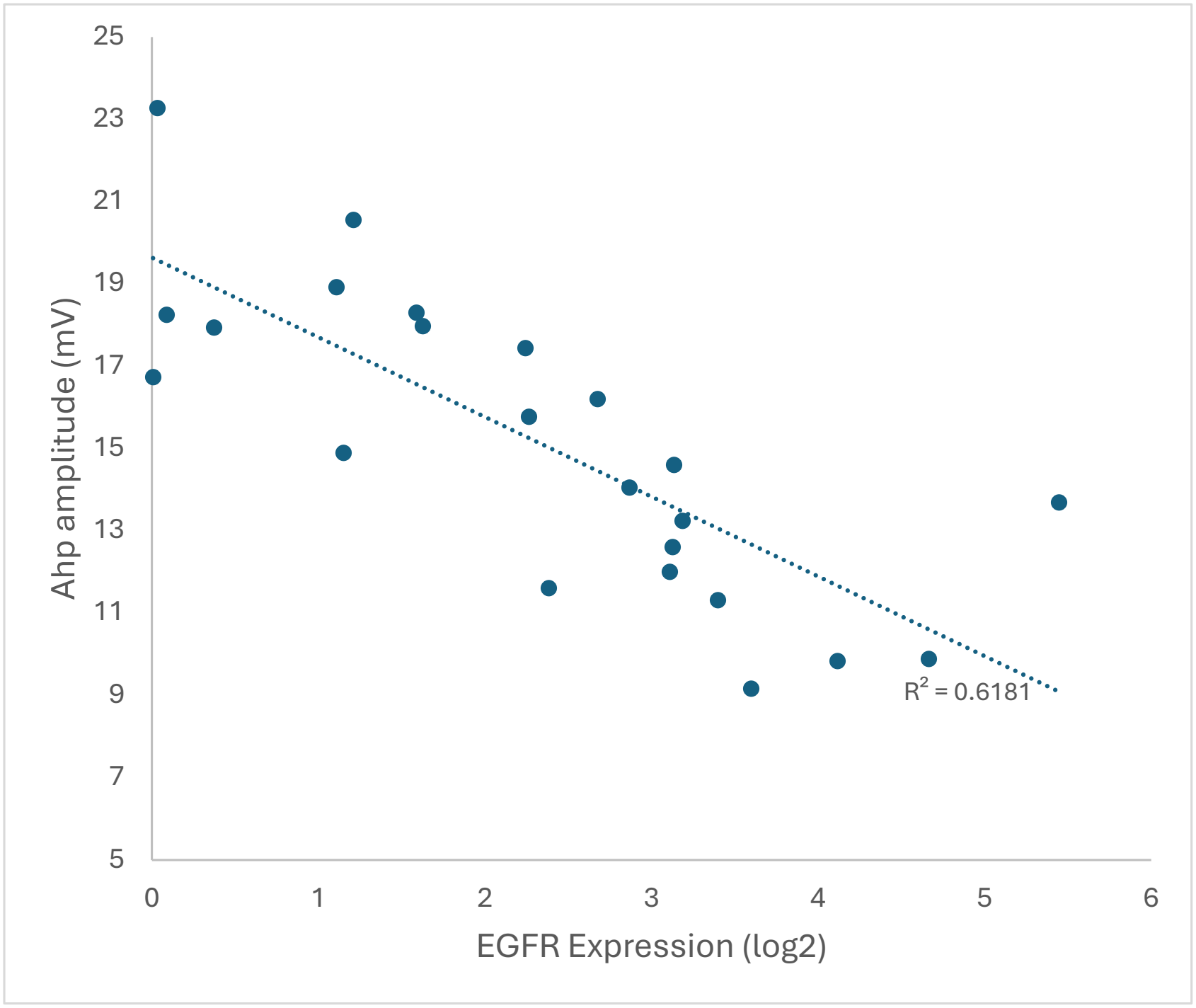
Anti-correlation between EGFR expression level and neuronal Ahp amplitude derived from Allen Brain Institute patch clamp and transcriptomic datasets using Enkefalos. This relationship is highly significant (*P*=8.71e^-6^). This relationship demonstrates that alterations of EGFR expression will affect neuronal electrophysiology.

## DISCUSSION

Omics technologies are rapidly expanding our view of the cellular events that contribute to disease progression and therapeutic treatment response. Given the volume of data produced by these methods, their full potential is highly dependent on the availability of bioinformatics tools for data interrogation and interpretation within the appropriate biological context. The study of neurological disorders requires the ability to link extensive transcriptomic data to altered neuronal function; however, there is currently a lack of bioinformatics tools to interpret lengthy lists of DE genes and proteins in the context of neuronal electrophysiology and associated cellular pathways. Here, we have developed a bioinformatics tool that fills this void. ENKEFALOS is easy to use, publicly available, and provides the ability to analyze lengthy lists of differentially expressed genes or proteins to identify those involved in alterations of neuronal electrophysiology and to further characterize their interactome networks and cellular pathways. Importantly, ENKEFALOS identifies key hubs among interactome networks. The identified key nodes are likely drivers of disease and potential therapeutic targets.

To demonstrate the utility of ENKEFALOS we performed a case study analysis of a list of differentially expressed genes from a seminal transcriptomic study of epilepsy in tuberous sclerosis complex (TSC). TSC is an autosomal dominant genetic disorder caused by mutations in the TSC1 and TSC2 genes, which code for hamartin and tuberin, respectively. Together these proteins inhibit the mammalian target of rapamycin (mTOR) pathway. Mutations in TSC1 and TSC2 prevent proper inhibition of mTOR, leading to uncontrolled cell growth and proliferation and the formation of benign hamartomas in several organ systems, including the brain. These lesions (tubers) are associated with epileptic foci and are sometimes surgically resected to treat drug resistant epilepsy. The molecular events downstream of mTOR that produce tubers and drive seizures in TSC are currently poorly understood. Disentangling the transcriptomic alterations associated with epileptogenesis, separately from lesion formation, is critical to understanding the molecular events that drive seizures and to help uncover new therapeutic targets.

In our presented case study of TSC data, ENKEFALOS was applied to the analysis of a list of genes that were significantly overexpressed in epileptic TSC tubers. ENKEFALOS provided several important results. First, the collective subset of DE genes having a significant correlation to neuronal electrophysiology were identified. Secondly, the interactome among this subset was characterized, including quantification of network connectivity. Highly connected key nodes identified likely drivers of epileptogenesis, and revealed the EGFR pathway as a pivotal node in TSC seizures. A growing body of evidence supports this finding and validates the results obtained with ENKEFALOS.

EGFR is a member of the c-erbB family of receptor tyrosine kinases, and primary ligands are epidermal growth factor (EGF) and transforming growth factor–α (TGFα) [16-18]. Stimulation of EGFR leads to activation of the ERK-MAPK and PI3K/Akt pathways and regulates a range of cellular functions [19]. While much EGFR research has focused on cancers, EGFR has also been implicated in TSC and epilepsy.

Significant overexpression of EGFR has been reported in cortical tubers of TSC patients and within the cortex of TSC mouse models [20, 21]. The study from which we obtained our demonstration set of genes also reported that EGFR signalling was among the most significantly enriched pathways [11]. Here, ENKEFALOS reveals that this pathway is significantly associated with alterations of neuronal electrophysiology, as opposed to formation of the neurological lesions (tubers) alone. Consistent with this finding, EGFR inhibitors were recently identified as potent anti-epileptic compounds in an epilepsy-on-a-chip system, suggesting the potential of utilizing EGFR inhibitors for alternative treatment option in TSC epilepsy [22].

Outside of TSC, EGFR overexpression has also been observed in other forms of epilepsy. A study of epileptic brain tissue isolated from patients with focal cortical dysplasia concluded that EGFR could be a novel biomarker of juvenile focal epilepsies [23]. A recent mega-analysis of somatic variants in 1386 individuals who underwent surgery for lesional epilepsies reported EGFR as a novel epilepsy risk gene in 3% of the cases [24]. These findings validate the value of using ENKEFALOS to mine omics datasets for key genes involved in altered electrophysiology.

In this work we present a web-accessible and easy-to-use bioinformatics tool to fill a critical void in neurological research that employs “omics” technologies. With the rapid pace of advancements in deep sequencing, single-cell sequencing, spatial transcriptomics, and proteomics, ENKEFALOS enables users to mine datasets for gene and protein expression events linked to altered neuronal electrophysiology. Furthermore, ENKEFALOS identifies the associated signalling pathways that converge on electrophysiological alterations and their key nodes, providing novel insight into disease mechanisms and potential therapeutic targets.

### Future Directions

The above case study demonstrates the ability of ENKEFALOS to identify key genes associated with altered electrophysiology within TSC. However, ENKEFALOS can function as a tool to identify genes and proteins associated with other forms of epilepsy, such as temporal lobe epilepsy, as well as other neurological disorders to identify signalling pathways and cellular networks that contribute to altered electrophysiology. ENKEFALOS can be continuously updated based on future discoveries, the use of different Cre-drivers to target different cell types and brain regions, and updates to the Allen Institute of Brain Sciences database. We anticipate that ENKEFALOS will be widely used in a broad range of neurological studies to provide new biological insights and hypotheses.

## Supporting information

Supplemental File 1 User Guide

Supplemental File 2 Electrophysiological definitions

Supplemental File 3 Case study input genes

## DATA AVAILABILITY

All code and data for Enkefalos are available at Github (https://github.com/MaayanLab/appyter-catalog/tree/main/appyters/ENKEFALOS_1/; https://github.com/MaayanLab/appyter-catalog/tree/main/appyters/ENKEFALOS_2).

The underlying data for transcriptomic correlates between gene expression and electrophysiological properties were downloaded from the GitHub site of the Pavlidis Lab as Online Tables 1-4 found here:

https://github.com/PavlidisLab/transcriptomic_correlates

## SUPPLEMENTAL MATERIAL

Supplemental File S1: User Guide

Supplemental File S2: Definitions of neuronal electrophysiological and morphological properties

Supplemental File S3: 3976 genes used as a case study

## ACKNOWLEDGMENTS

Gratitude is expressed for the work of the Pavlidis laboratory at the University of British Columbia which created the initial assembly of the Allen Brain data into a database and laid the foundation for Enkefalos.

Publication of Enkefalos as an Appyter application was made possible by the Ma’ayan laboratory at the Icahn School of Medicine.

## AUTHOR CONTRIBUTIONS

Krish Upadhyay: Methodology, Software, Data Curation, Writing. Dylan Ukasik: Conceptualization, Investigation, Writing. Douglas Craig: Software. Alan Dombkowski: Supervision, Conceptualization, Investigation, Writing.

