## Supplemental File 1 User Guide for "ENKEFALOS: Electrophysiological NetworKs Elicited From ALtered OmicS"

### Εγκέφαλος

*User's Guide*

**Version 1.0**

Krish Upadhyay
Dylan Ukasik
Douglas Craig
Alan Dombkowski

**Wayne State University**
School of Medicine
Department of Pediatrics
Dombkowski Lab

### Introduction

ENKEFALOS (Electrophysiological NetworKs Elicited From ALtered OmicS) is a tool designed to help identify genes that are significantly associated with neural electrophysiological and morphological measures. Electrophysiological and morphological data from a wide variety of neural cell types were taken from the Allen Brain Institute and computationally correlated to gene expression measures by a Pavlidis lab study. These correlatory measures were then organized into several files which ENKEFALOS tabulates through based on the user's inputs. Rather than focus on single gene-measure correlations, ENKEFALOS takes a list of genes and is able to identify genes with significant correlations given a user-defined FDR (q-value) threshold. ENKEFALOS is able to do more downstream analysis using StringDB to identify known interactions between genes, the number of interactions each of these genes have, and potential cellular drivers of the user's disease of interest. A secondary ENKEFALOS application exists for more specific gene-measure correlation analyses, including correlation plots, plot data, and a subset StringDB interactome with the user's gene of interest as the central node. For the primary ENKEFALOS application, the input consists of a list of genes (manually inputted or a .csv file uploaded) and a FDR threshold, with the output being downloadable .csv files and a link to the StringDB analyses. The secondary ENKEFALOS application requires an input of a gene of interest, an electrophysiological/morphological measure, and the list of enriched genes from the primary analysis (manually inputted or a .csv file uploaded). The outputs for the secondary application consist of downloadable .csv files with the results and a link to the subset StringDB network. This software was written by Krish Upadhyay under the mentorship of Dr. Alan Dombkowski and with the resources of the Dombkowski lab. ENKEFLAOS's results have successfully demonstrated its validity on several datasets of differentially expressed genes from a wide variety of neurological disorders, including TSC epilepsy.

### Getting Started

---

##### 1. Input/Upload Gene List and Input FDR Threshold

You may try using the sample list of genes or use your own genes of interest from your study.

##### 2. Run Primary Analysis

Click on the submit button at the bottom of the page. Be sure all fields are filled.

##### 3. Evaluate and Save Primary Results

Inspect the results. For results involving tables, you will be provided a downloadable .csv file. For the StringDB analysis, you will be provided the link.

##### 4. Input Gene and Property Measure for Single Gene Analyses

Preferably, use a gene-property pair of interest from your primary analysis. You will also have to input your enriched genes from the first analysis for the subset network.

##### 5. Run Secondary Analysis

Click on the submit button at the bottom of the page. Be sure all fields are filled.

##### 6. Evaluate and Save Secondary Results

Inspect the results. For results involving tables, you will be provided a downloadable .csv file. For the StringDB subset analysis, you will be provided the link. For the plot analysis, you will be able to save the image.

### Loading and Inputting Data

---

The ENKEFALOS analysis begins with the primary Appyter. You will have to submit a gene list, which can be done in two manners, and input a FDR (q-value) threshold.

#### ***Primary Analysis***

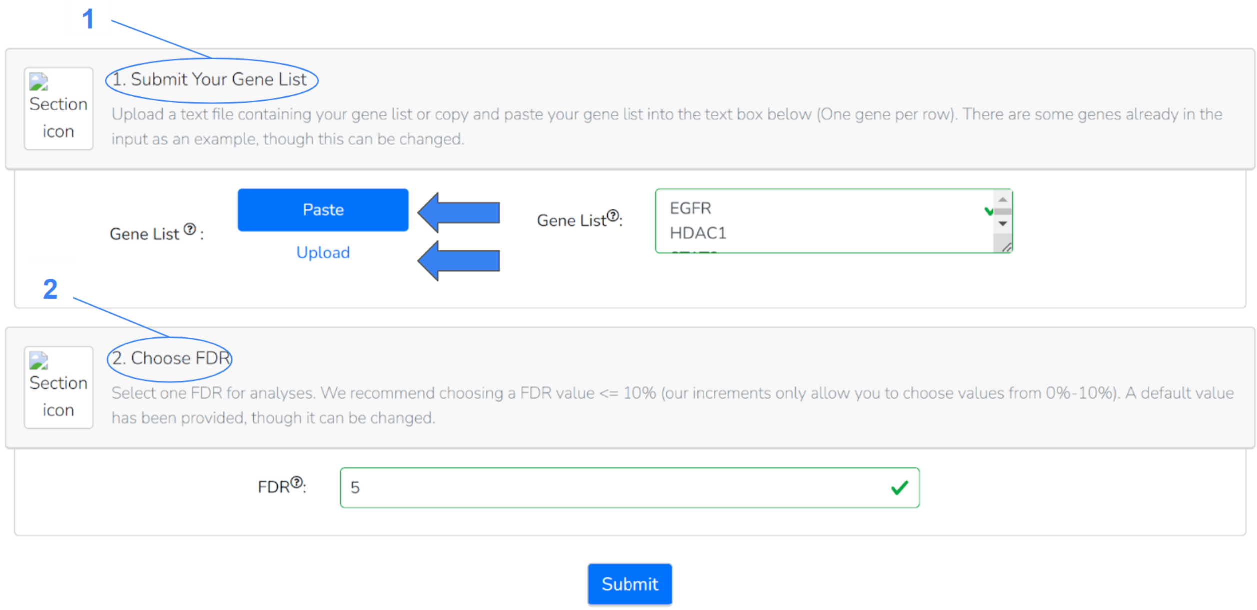


**Figure 1.** Inputting a gene list and FDR threshold

##### 1. Submitting a Gene List

- A gene list may be inputted by one of two ways: pasting a list separated by spaces, tabs, or lines, or uploading a .csv file containing your genes of interest.
- Clicking on one of the two options specified by the arrows above will allow you to choose your mode of input.
- By default, an example gene list has been provided to you that you may run for testing purposes.

##### 2. Choosing a FDR Threshold

- Type in a FDR (q-value) threshold for your analysis. You are allowed to choose a value from 0%-10% (0-0.10). We recommend using a FDR of 10% or less for significant results.
- By default, a value of 5% has been inputted, though this can be changed manually or by using the scroll bar on the right side of the input field.

#### *Secondary Analysis*

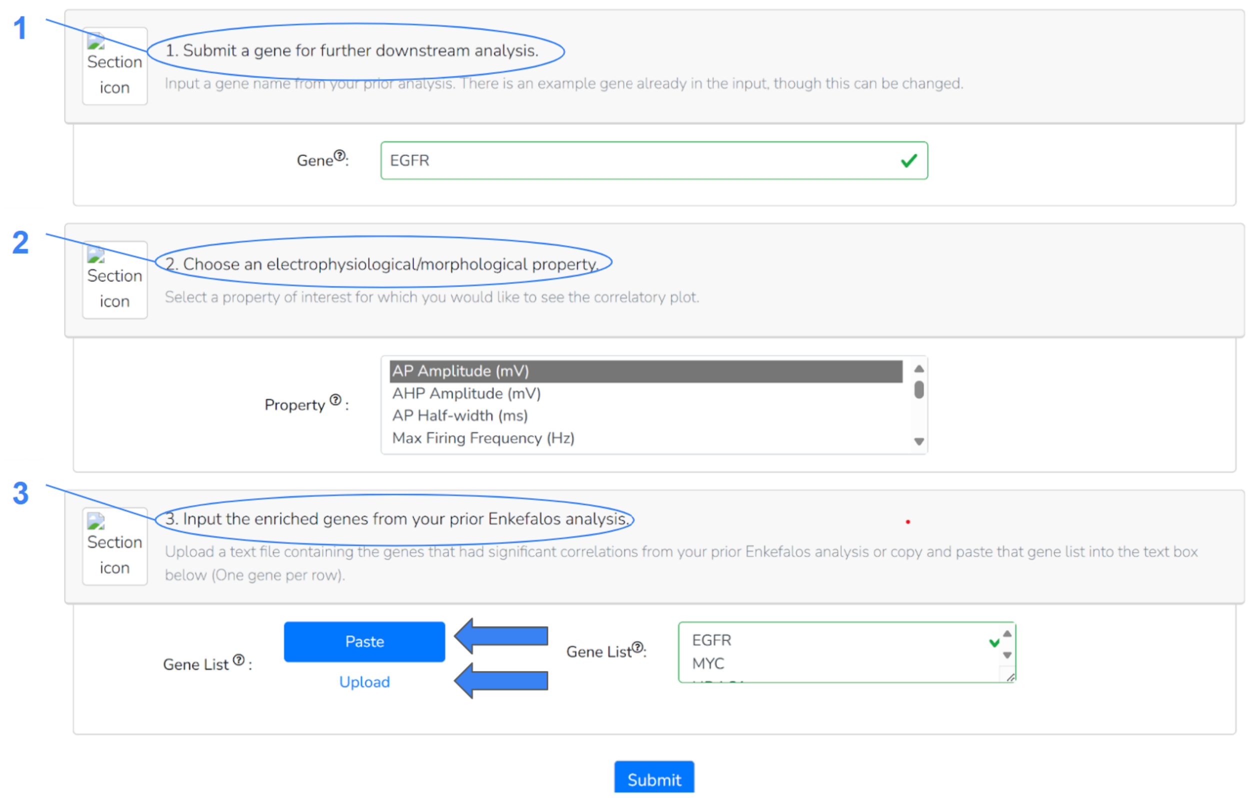


**Figure 2.** Inputting a gene of interest, property measure, and enriched gene list from primary analysis

##### 1. Inputting a Gene of Interest from Prior Analysis

- A gene of interest from your prior analysis may be inputted here for further downstream analysis.
- By default, an example gene has been provided to you that you may run for testing purposes.

##### 2. Choosing a Property Measure of Interest

- Select a property measure of interest that you would like to see correlated with your gene of interest.
- By default, action potential amplitude (mV) has been selected.

##### 3. Submitting List of Enriched Genes from Prior Analysis

- Either paste a list or upload a .csv file containing the enriched genes (genes that had significant correlations to properties) from your prior analysis to form a subset network with your gene of interest as the central node.
- You may directly upload the resultant genes with significant interactions .csv file (Significant_Interactions) from your prior analysis into this field.

### Evaluating and Saving Results

After submitting your inputs for both ENKEFALOS appyters, there are three categories of information from which you may assess your results:

- Tables including a variety of analysis information
- StringDB gene interactomes
- Correlation Plots

Each is available under a separate analysis section in the Appyters and is described in detail below.

#### ***Primary Analysis***

##### 1. Significant Correlations

| gene_symbol | property | FDR_gene\|class_anova |
|---|---|---|
| EGFR | ahpamp | 0.000003 |
| EGFR | aphw | 0.000006 |
| EGFR | f_i_curve_slope | 0.000087 |
| EGFR | maxfreq | 0.000243 |
| MYC | ahpamp | 0.001413 |
| MYC | f_i_curve_slope | 0.006255 |
| EGFR | E_PC1 | 0.006441 |
| EGFR | avg_isi | 0.008331 |
| MYC | rheo | 0.016571 |
| EGFR | cap | 0.019936 |

Download CSV file of the full table of significant results

**Figure 3.** Significant correlations result from primary analysis

- **gene_symbol**: Genes with significant correlation to associated property in the table based on the Pavlidis lab study.
- **property:** Property measure with significant correlation to associated gene in the table based on the Pavlidis lab study.
- **FDR_gene|class_anova:** False discovery rate (q-value) of gene-property correlation. All will be under user-specified threshold.

##### 1. Significant Correlations (cont.)

Although only the top ten significant gene-property correlations are shown, you may download the full .csv table by clicking the link specified by the arrow in the image above.

#### 2. Genes with Significant Interactions

This analysis provides you with the list of enriched genes from your analysis (genes that had significant correlations with property measures given your specified FDR threshold). The result will be formatted exactly the same as the prior analysis; a table with the top ten genes with correlations and a downloadable .csv file to the full list. <u>You will use this list as an input if you choose to do the secondary analysis.</u>

#### 3. String Call and Network Analysis

Formatted the same as the prior two tables, you will be provided a table with genes from your enriched genes list that had one or more known interactions determined by the StringDB application. Theoretically, genes with the most interactions may be potential cellular drivers of your disease of interest. A link to the StringDB network will be provided too if it does not pop up.

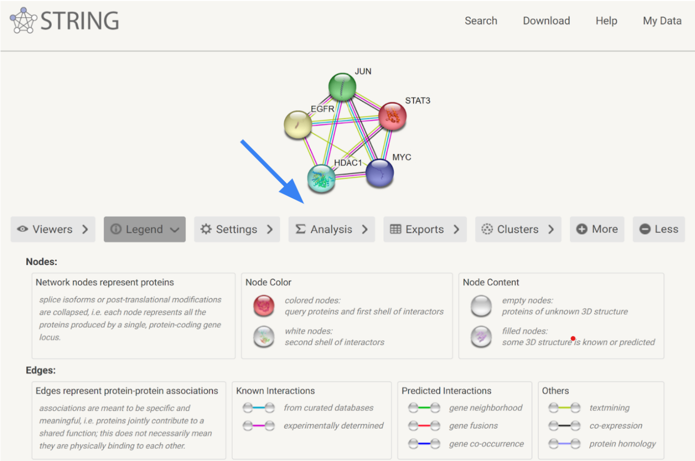


**Figure 4.** StringDB call with list of enriched genes.

- **Analysis:** Specified by the arrow, this tab will allow you to view network statistics and determine cellular and biological processes that genes in your network are involved in.
- Other tabs are there to change the settings of your interactome, view more information regarding it, or export its information.

### ***Secondary Analysis***

#### 1. Correlation Graph

This is a correlation plot between your gene of interest and chosen property measure. Your gene of interest's count per million in a given neural cell type (done by the Allen Brain institute) is log2-transformed and added to 1 as a measure of gene expression and plotted against the x-axis. The chosen property's measure (also done by the Allen Brain institute) is plotted against the y-axis. You may copy/save this image directly from the ENKEFALOS application for your desired correlation.

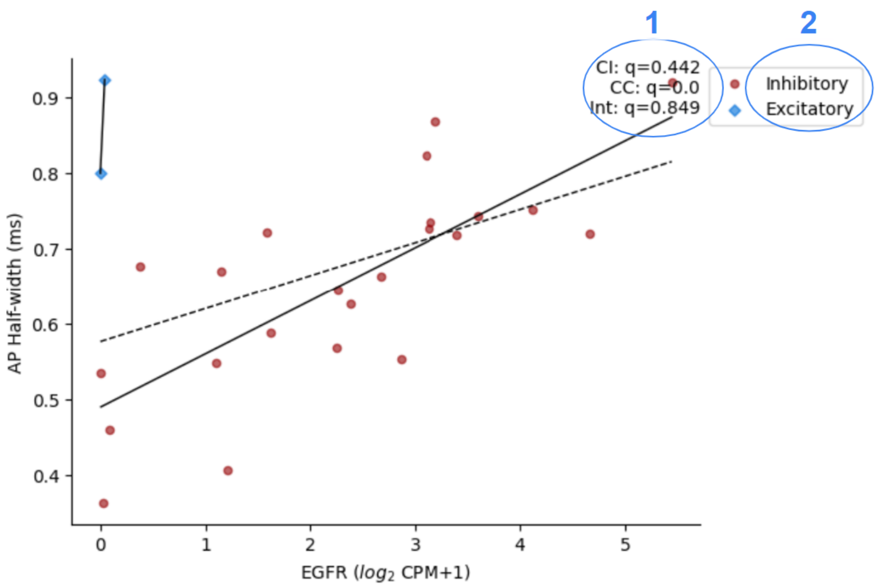


**Figure 5.** Example correlation plot of EGFR and action potential half-width.

- **CI** (Class-independent), **CC** (Class-conditional), **Int** (Interaction): Methods used by the Pavlidis lab study to model gene-property relationships. More information about them can be found in their study. ENKEFALOS uses the CC model for significance (FDR/q-value). The solid lines represent gene-property relationships, while the dashed line represents a cross-class difference.
- **Inhibitory** (Inhibitory neuron), **Excitatory** (Excitatory neuron)

#### 2. Plot Data

For your convenience, this analysis section allows you to download the .csv files of both the excitatory and inhibitory data plotted on your correlation plot of interest.

| | A | B | C |
|---|---|---|---|
| 1 | | AP Half-width (ms) | EGFR |
| 2 | Gad2-IRES-Cre__L5__inh | 0.405714286 | 1.211382873 |
| 3 | Sst-IRES-Cre__L6__inh | 0.499375 | 0 |
| 4 | Nos1-CreERT2\|Sst-IRES-FlpO__L5-L6__inh | 0.755428571 | 0 |
| 5 | Oxtr-T2A-Cre__L5-L6__inh | 0.676666667 | 0.375397526 |
| 6 | Chrna2-Cre_OE25__L5__inh | 0.583448276 | 0 |
| 7 | Chrna2-Cre_OE25\|Pvalb-T2A-Dre__L5-L6__inh | 0.46 | 0 |
| 8 | Chat-IRES-Cre-neo__L2/3__inh | 0.743555556 | 3.596237233 |
| 9 | Htr3a-Cre_NO152__L4__inh | 0.55225 | 2.86865861 |
| 10 | Sst-IRES-Cre__L4__inh | 0.433478261 | 0 |
| 11 | Pvalb-IRES-Cre__L4__inh | 0.359206349 | 0 |
| 12 | Htr3a-Cre_NO152\|Pvalb-T2A-Dre__L5-L6__inh | 0.418 | 0 |
| 13 | Vip-IRES-Cre__L6__inh | 0.726551724 | 3.126052009 |
| 14 | Vip-IRES-Cre__L5__inh | 0.717647059 | 3.398294243 |
| 15 | Ndnf-IRES2-dgCre__L1__inh | 0.919487179 | 5.44489703 |
| 16 | Chrna2-Cre_OE25__L6__inh | 0.548 | 1.106736494 |
| 17 | Htr3a-Cre_NO152__L2/3__inh | 0.735681818 | 3.136392281 |
| 18 | Nos1-CreERT2__L1-L4__inh | 0.8675 | 3.186680345 |
| 19 | Pvalb-IRES-Cre__L6__inh | 0.357368421 | 0 |
| 20 | Vip-IRES-Cre__L4__inh | 0.75125 | 4.116467755 |
| 21 | Pvalb-IRES-Cre__L5__inh | 0.360855263 | 0 |
| 22 | Nkx2-1-CreERT2__L5-L6__inh | 0.721538462 | 1.58732587 |
| 23 | Chat-IRES-Cre-neo__L4__inh | 0.72 | 4.663961955 |
| 24 | Gad2-IRES-Cre__L1__inh | 0.822857143 | 3.111289552 |
| 25 | Vip-IRES-Cre__L1-L2/3__inh | 0.62625 | 2.383545824 |
| 26 | Vipr2-IRES2-Cre__L1-L6__inh | 0.567272727 | 2.244872923 |

**Figure 6.** Inhibitory correlation plot data for EGFR-AP Half Width correlation.

- **A (Cell Type):** Inhibitory or excitatory depending on which file you download. How cell types were differentiated and other details may be found in the Pavlidis Lab study.
- **B (Property Measure):** All the measured values of the given property from the associated cell types in the table.
- **C (Gene Expression):** All the gene expression measures of the given gene from the associated cell types in the table.

#### 3. Subset Network

Provides you with another StringDB network with your gene of interest as the central node of the enriched gene interactome (all genes directly connected to your gene of interest will be in this network). This provides the user a way to more closely investigate which genes from the list of enriched genes their gene of interest may be interacting with.

### Contact Us

Please contact us with any issues, thoughts, questions or concerns about ENKEFALOS. We would appreciate any insights regarding the application.
