## Supplemental File 2 Electrophysiological definitions for "ENKEFALOS: Electrophysiological NetworKs Elicited From ALtered OmicS"

| Property | Definition |
|---|---|
| Soma Surface | Surface area of the cell body |
| Total Volume | Volume of the cell, including cell body as well as processes |
| Total Length | Total length of all processes |
| Max Branch Order | Maximum number of times that a process bifurcates between the soma and branch tip |
| Branchiness | Number of bifurcations encountered per process length |
| Bifurcation Angle | Mean angle across all bifurcation points |
| Morphology PC1 (44%) | N/A |
| Morphology PC2 (21%) | N/A |
| Morphology PC3 (19%) | N/A |
| Adaptation Ratio | Ratio of durations between early and late AP inter-spike intervals in an AP train |
| Input-Output Curve Slope | Slope of the relationship between current injection and resulting firing frequency, based on multiple long current steps |
| Max Firing Frequency | Maximum observed AP discharge rate |
| Latency | Latency to fire the first action potential during a long current step |
| ISI CoV | Variability between interspike intervals within one sweep, measured as standard deviation/mean |
| Average Interspike Interval | Average time elapsed between spikes during a sweep |
| Sag | Measure of the extent to which the membrane potential recovers toward resting potential when the neuron is strongly hyperpolarized (between -90 and -110 mV) |
| Resting Membrane Potential | Membrane potential at the onset of whole-cell recording |
| Input Resistance | Input resistance measured at steady-state voltage response to current injection |
| Time Constant Tau | Time constant for the membrane to repolarize after a small current injection of fixed amplitude and duration |
| Capacitance | Neuron capacitance, typically measured by dividing membrane time constant by membrane resistance |
| AHP Amplitude | Calculated as the voltage difference between AP threshold and AP trough. Commonly defined using first AP in train at rheobase current. |
| AP Amplitude | Voltage indicating height of action potential. Usually calculated as the difference between AP peak and AP threshold voltages. Commonly measured using first AP in train at rheobase current. |
| AP Half-width | Calculated as the AP duration at the membrane voltage halfway between AP threshold and AP peak. Most commonly calculated using first AP in train at rheobase current. |
| AP Threshold | Voltage at which AP is initiated (as assessed by measuring rising slope of membrane voltage) |
| Rheobase | Minimum current injected somatically required to fire AP |
| Electrophysiology PC1 (47%) | N/A |
| Electrophysiology PC2 (18%) | N/A |
| Electrophysiology PC3 (9%) | N/A |
